# Life on the margin: rainwater tanks facilitate overwintering of the dengue vector, *Aedes aegypti*, in a sub-tropical climate

**DOI:** 10.1101/517318

**Authors:** Brendan J Trewin, Jonathan M Darbro, Myron P Zalucki, Cassie C Jansen, Nancy A Schellhorn, Gregor J Devine

## Abstract

A key determinant of insect persistence in marginal habitats is the ability to tolerate environmental extremes such as temperature. *Aedes aegypti* is highly invasive and little is known about the physiological sensitivity of the species to fluctuating temperature regimes at the lower critical threshold. This has implications that limit establishment and persistence of the species in sub-optimal regions. Daily winter temperatures were measured in common Australian larval habitats, replicated in environmental chambers, and used to investigate the effect of fluctuating temperatures on the development and survival of tropical and subtropical strains of Australian *Ae. aegypti*. Development was slow for all treatments but both strains were able to complete development to the adult stage, suggesting previous models underestimate the potential for the species to persist in eastern Australia. Results suggested that thermal buffering in large volume habitats, and water that persists for greater than 32 days, will facilitate completion of the life cycle during sub-tropical winters. Furthermore, we provide a non-linear estimate of the lower critical temperature of *Ae. aegypti* that suggests the current threshold may be incorrect. Our study demonstrates that the current re-introduction of water storage containers such as rainwater tanks, into major Australian population centres will increase the risk of *Ae. aegypti* establishment by permitting year-round development south of its current distribution.

## Introduction

A key determinant of insect distribution and persistence is the ability of a species to tolerate micro-climates at a local scale (1). Conditions within the core distribution of a species will be near-optimal and less stable populations will persist around the margins of an insect’s distribution; the permanency mediated by access to food sources, the availability of suitable oviposition and resting sites and the abiotic factors associated with these micro-habitats (2-4). In recent years there has been renewed interest in predicting the spread of the mosquito *Aedes aegypti* (L.) into cool range margins, primarily due the increased variability in temperature and rainfall associated with climate change and the importance of the species as a disease vector (4). In particular, rising temperatures, unpredictable rainfall and urban landscapes that are evolving in response to climatic changes may impact mosquito distributions, daily activity patterns and peak annual population abundance in marginal habitats (2, 3, 5-8).

*Aedes aegypti* is a highly anthropophilic species (2, 9). The continuous availability of oviposition sites and blood meals afforded by intra-domiciliary habitats can mitigate otherwise hostile environments and has allowed the species to achieve a global distribution (10, 11). Pertinent to the spread and re-establishment of *Ae. aegypti* in parts of Australia and other marginal habitats is the increasing presence of large permanent water storage containers, namely domestic rainwater tanks. The temperature-buffering effect of these tanks may allow continual *Ae. aegypti* development in marginal habitats (2). Water has a high specific heat capacity, low thermal inertia and, in large volumes, can resist large and rapid fluctuations in temperature (12). These tanks can also provide permanent aquatic habitats throughout the year. It is hypothesised that the presence and then removal of rainwater tanks may have contributed to historical patterns of *Ae. aegypti* distribution across temperate Australia (3). We believe that the modern trend for the widespread installation of large water storage containers, in response to an unpredictable climate, may increase the risk of re-establishment and expansion.

The lower critical temperature for the development of larval stages of *Ae. aegypti* is widely accepted to be approximately 11.8°C (4, 13). Methodologies to estimate thermal performance rely primarily on observations of development at constant temperatures and the use of linear regression to estimate lower critical thresholds (4). Due to the difficulty of working at extremes in the thermal performance curve, little research has defined the lower critical threshold of *Ae. aegypti* using non-linear methods.

It has been recognized for some time that insects subjected to constant temperatures in the laboratory do not accurately reflect development and survival in the field (14). Studies using fluctuations in temperature more accurately reflect the natural daily cycles experienced by insects. In mosquitoes, fluctuations in water temperature can alter immature mosquito development and survival around critical development temperatures. Studies suggest that thermal tolerance is improved when fluctuating temperature regimes are compared with those using constant temperatures (13, 15, 16). For instance, Carrington et al. (13) found that a large diurnal thermal range of 18.6°C around a mean of 16°C significantly reduced development time (but not survival) of *Ae. aegypti* when compared to small (7.6°C) fluctuations or constant temperatures. There has been little research on the survival and development of *Ae. aegypti* in fluctuating temperatures and their relation to potential geographic distribution, particularly around the lower critical threshold.

To examine whether rainwater tanks encourage the survival and development of *Ae. aegypti* under temperate Australian conditions, the abiotic conditions typical of tanks (limited temperature fluctuations) and smaller containers (high fluctuations) were measured during winter in Brisbane, Australia. Those fluctuating temperatures were then replicated in environmental chambers and we assessed their impact on the larval development and survival of a tropical and subtropical strain of Australian *Ae. aegypti*. It was hypothesised that 1) the temperature buffering provided by rainwater tanks will increase survival and time to adult emergence when compared to smaller volume habitats and 2) that the subtropical *Ae. aegypti* population would have higher survival and faster development under winter conditions than *Ae. aegypti* sourced from the tropics, due to adaptations to local conditions.

## Methods

### Environmental observations

Temperature fluctuations in different container types were measured over winter in Brisbane (27.47° S, 153.03° E), from the start of June until September, 2014. HOBO Pendant®Data Loggers were placed into ten rainwater tanks and ten buckets, and tank locations and classified as high (>66%), medium (33-65%) and low (0-32%) shade categories. Shade categories were estimated visually as the percentage of structural or vegetative shade covering each container. Air temperatures were measured outside in a high shade location. Tanks were checked fortnightly to ensure they remained sealed to the ingress of adult mosquitoes. Locations of tanks included: Greenslopes (27.51°S, 153.05°E), Moorooka (27.54°S, 153.03°E), Salisbury-Nathan (27.55°S, 153.03°E), Sunnybank (27.58°S, 153.06°E), Camira-Gailes (27.63°S, 152.91°E), Indooroopilly (27.50°S, 152.97°E) and St Lucia (27.50°S, 153.00°E; Fig 1). A black 9L bucket, representing the most common type of container in Brisbane backyards (Darbro, pers. comm.), was placed under similar shade conditions in a northerly position next to each tank.

**Fig 1.**
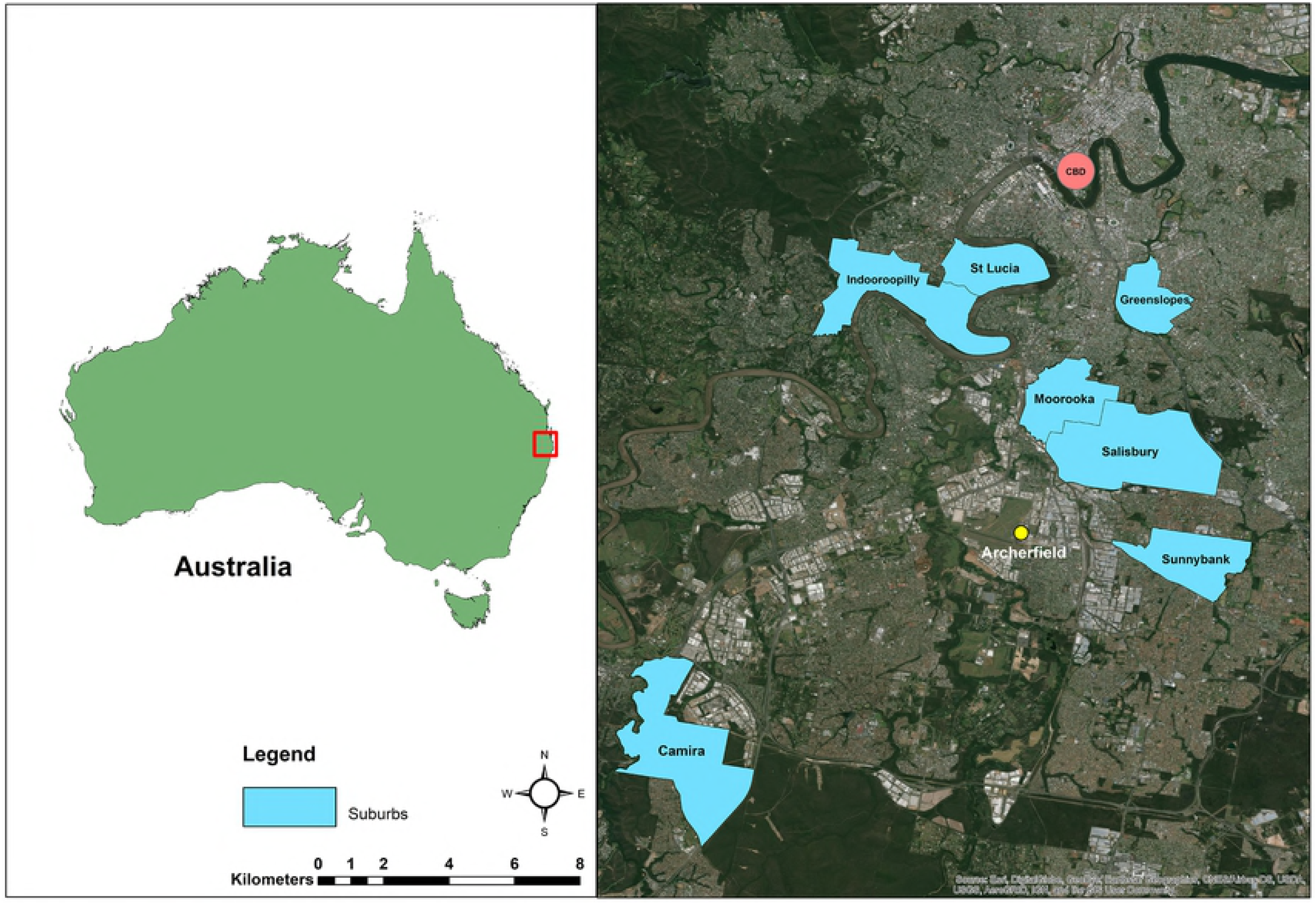
Location of study sites in Brisbane, Australia. Suburbs where temperatures in container habitats were measured during winter, 2014. Climate data was taken from Archerfield (yellow circle)(17). Map Source: Base layer of Brisbane region sourced from Esri World Imagery © (18); Esri, DigitalGlobe, GeoEye, Earthstar Geographics, CNES/Airbus DS, USDA, USGS, AeroGRID, IGN, and the GIS User Community

Data loggers were attached to a floatation device in each tank, submerged to a depth of 30cm below the water surface to avoid surface temperature fluctuations. Flotation devices were attached to a tape measure suspended from the roof of the tank. Water losses due to evaporation were noted fortnightly from tanks and buckets. Ten tank abiotic characteristics were measured including temperature inside and out of each tank, total volume of water, humidity, dew point, pH, salinity, conductivity, total dissolved solids, and presence of larvae inside tanks and “first flush” devices (a separate pipe for collecting sediment before water enters a tank). The five-sweep netting technique was used to sample mosquito larvae from rainwater tanks (19). As *Ae. aegypti* is not currently present in Brisbane, the native tree hole mosquito, *Aedes notoscriptus*, was used as an indicator species for tanks productivity. After the first survey, two tanks were disconnected from input water sources to estimate evaporation rate and to prevent ingress of mosquito larvae and eggs. Archerfield (−27.57°S, 153.01°E) climate data was selected as it is the closest Bureau of Meteorology (17) station to the rainwater tanks surveyed.

### Survival and development rate trial

*Aedes aegypti* colonies were established from eggs sourced from field sites in Cairns (tropical strain; −16.92°S, 145.78°E) and Gin Gin (subtropical strain; −24.99°S, 151.95°E), Queensland, in January, 2015. The subtropical colony originated from 30 eggs collected from ovitraps at three separate houses, while the tropical colony was established from >100 eggs collected from a single ovitrap at five separate properties. North Queensland is within the optimal range of the species and has the highest genetic diversity of Australian populations (20). A PCR test for the presence of *Wolbachia* (21) revealed absence in both colonies (n=60). Colonies were maintained at QIMR Berghofer at >500 individuals per generation, and insectary conditions held at 26 ± 1°C, a 70% (± 10%) relative humidity, with a 12:12 hour light cycle with twilight period. Adults were blood fed on an adult volunteer for 15 minutes, two days after emergence for three consecutive days (QIMR Berghofer Medical Research Institute human ethics form P2273). Eggs were collected from both colonies after generation two for use in environmental chamber experiments. Eggs were hatched synchronously using vacuum immersion in water at room temperature (24°C) for one hour.

A fluctuating temperature regime was derived from tank and bucket measurements during the coldest week in Brisbane during July 2014. These fluctuations were replicated in environmental chambers using two hourly intervals (S1 Appendix). A control treatment was set at 26°C (± 1°C), 70% (± 10%) Light regimes for all larval treatments were set at a 10:14h cycle, typical of Brisbane in July. Humidity for environmental chambers was set at 75% which is comparable to those observed in rainwater tanks during winter (S2 Appendix).

Fifty first-instar larvae from each mosquito strain were transferred into each of eight white, plastic containers (183 × 152 × 65mm) for a total of 400 larvae per strain, per temperature treatment, in a randomized block design and 500mL of appropriately chilled tap water (aged 2 days) was placed into each container. Larvae were fed with TetraMin^©^ground fish food (Tetra, Germany) standardized to the high diet treatment of Hugo et al. (22), with food concentrations estimated per larvae per volume each day and excess food removed daily before feeding. Containers were topped up daily with chilled water. Trays were rotated within environmental chambers and insectary shelves daily to prevent location bias. Containers were photographed each day to facilitate counting of all surviving instars and adult emergence was also recorded (defined as complete emergence from the pupal case).

### Statistical analysis

To assess the effect of temperature on survival to adult in all treatments, Kaplan Meier (log-rank) survival analysis was used (23). Student’s t-tests and ANOVA were used to compare mean survival, development times and degree days for *Ae. aegypti* strains in tanks, buckets and controls. Student’s t-tests were used to compare air temperature, humidity and dew point from measurements taken inside and outside the tanks. Heating degree day (HDD) models were constructed at 30 minute intervals, with a lower critical temperature of 11.78°C for the constant, tank and bucket temperature treatments. Statistical significance between each HDD model was compared with t-tests. For an estimate of cold stress, a cooling degree day (CDD) model was calculated for bucket treatments. All analyses were done using R version 3.2.2 (24) with the ‘nlme’, ‘survival’ libraries and ‘survminer’ used for plotting survival curves. Map of Brisbane suburbs was created using ArcGIS® 10.5 software by Esri (ESRI® Inc., Redlands, CA, USA).

## Results

### Field Observations

During winter 2014 Brisbane experienced average rainfall conditions, with above average maximum air temperatures, below average minimum temperatures and a minimum temperature of 0.5°C (17; S3 Appendix). Rainwater tanks had a mean temperatures of 16.8°C (range = 11.3°C, SD = 1.9) while buckets had a mean temperature of 16.3°C (range = 29.9°C, SD = 4.1; S2 Appendix). The relative difference between the mean weekly temperature in tanks consistently stayed above the mean weekly air temperature throughout the winter (mean relative difference = 1.3°C, SD = 0.14), while the relative difference in buckets was 0.55°C (SD = 0.14; Fig 2). The mean hourly tank temperature in high and low shade did not drop below the lower critical temperature during July (Fig 3). The minimum temperature of tanks only dropped below the lower critical temperature on 5.4% (5 of 92 days) (Fig 4). Temperatures below the lower critical temperature coincided with tanks in high shade or containing under 500L of water at the time of measurement (Fig 4, S2 Appendix).

**Fig 2.**
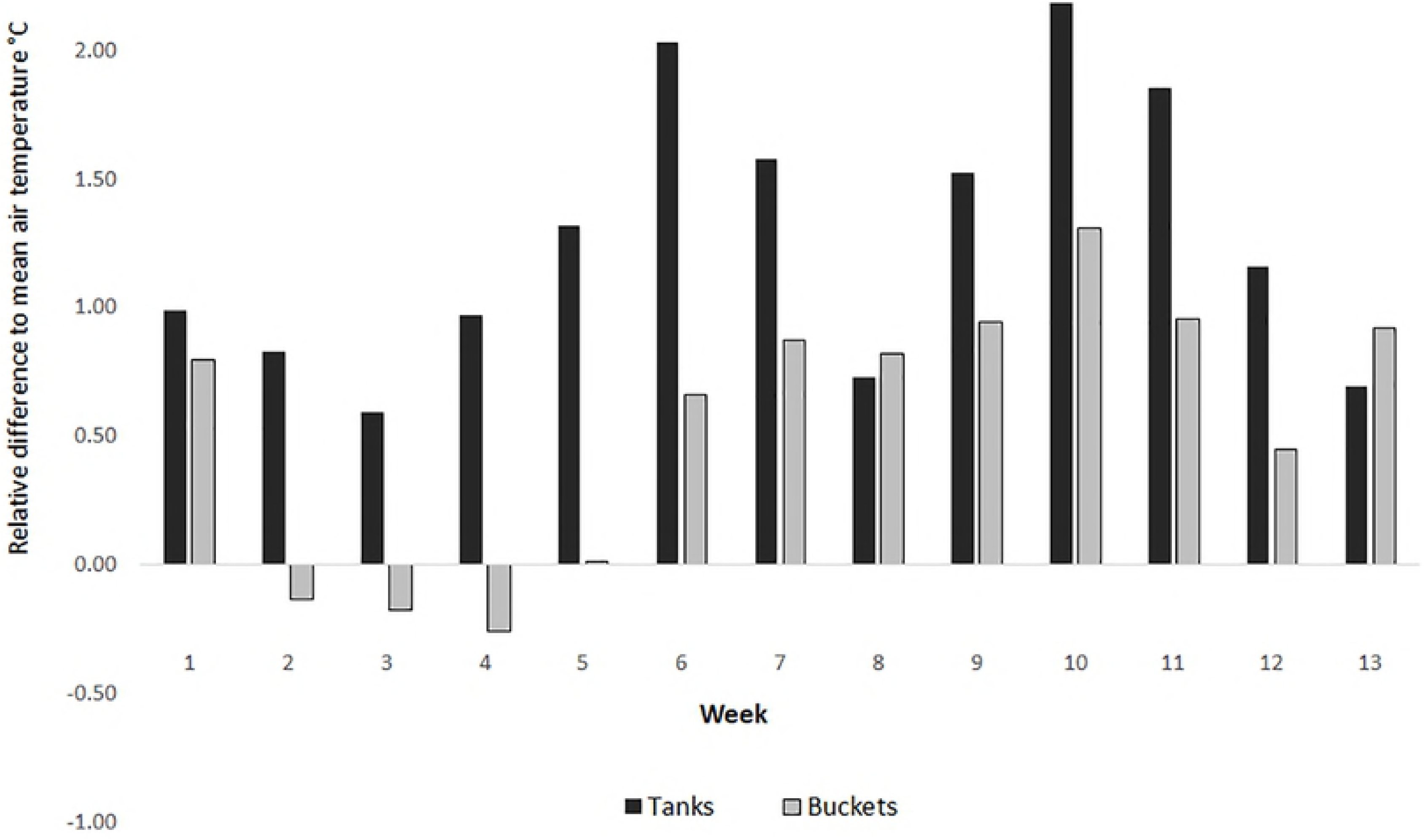
Relative differences between weekly container water and air temperatures. Relative difference in mean weekly water temperature for rainwater tanks and buckets to air temperature in 100% shade during winter in Brisbane, 2014.

**Fig 3.**
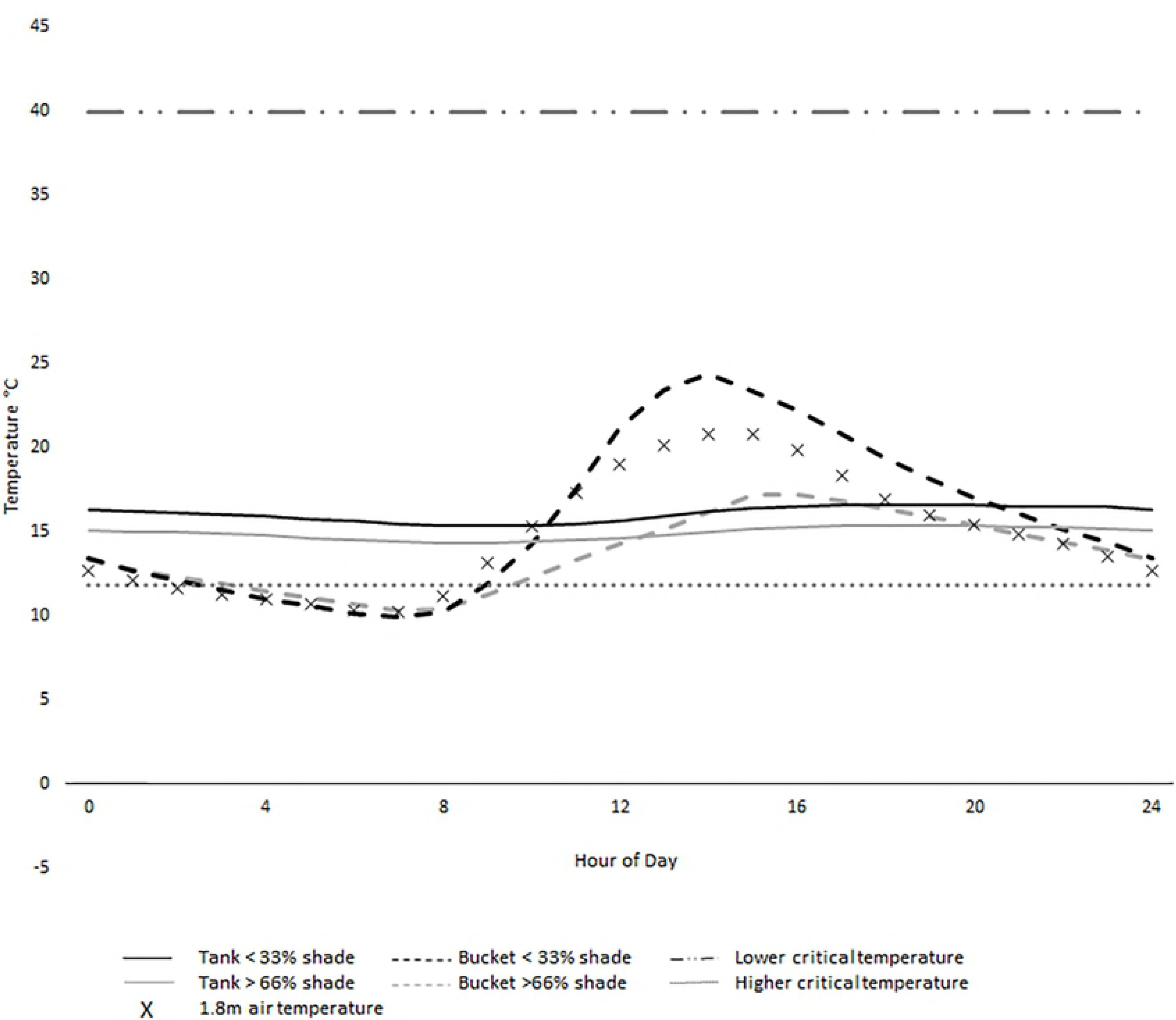
Mean hourly water temperatures from buckets and rainwater tanks during July in Brisbane, 2014. Air temperatures are recorded from 100% shade (crosses) and critical thresholds of *Aedes aegypti* are displayed.

**Fig 4.**
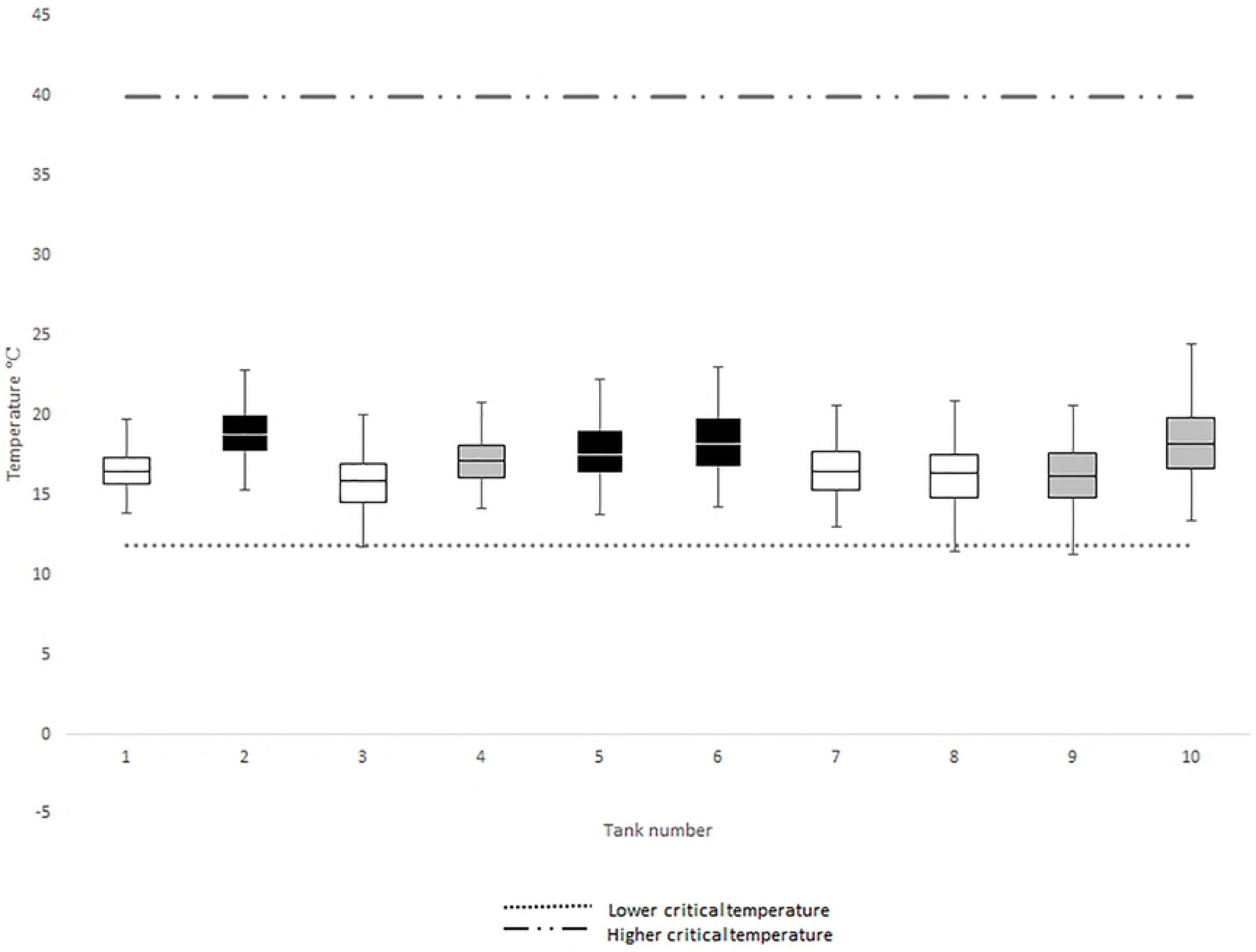
Interquartile ranges of water temperatures of individual rainwater tanks in Brisbane during winter 2014. Lower critical temperature threshold (11.8°C) for *Aedes aegypti* taken from Eisen et al. (4) and upper threshold (40.0°C) from Richardson et al. (39). Black, grey and white boxes represent low, moderate and high shade conditions, respectively.

In buckets the mean hourly temperature (high and low shade regimes) and mean daily minimum for all shade regimes dropped below the lower critical temperature throughout July (Fig 3) and all months during winter (Fig 5), respectively. Daily temperatures in buckets dropped below the lower critical temperature 66.3% (61/92) and 93.3% (28/30) of the time in winter and July, respectively. During July, the lowest temperatures observed in buckets, tanks and air was 5.4°C, 11.2°C and 0.5°C respectively. Differences between mean internal (21.4°C, SD = 0.99) and external (21.7°C, SD = 0.94) air temperatures of tanks at different shade levels measured fortnightly were not significant (F(1,12) = 0.26, *P* = 0.80).

**Fig 5.**
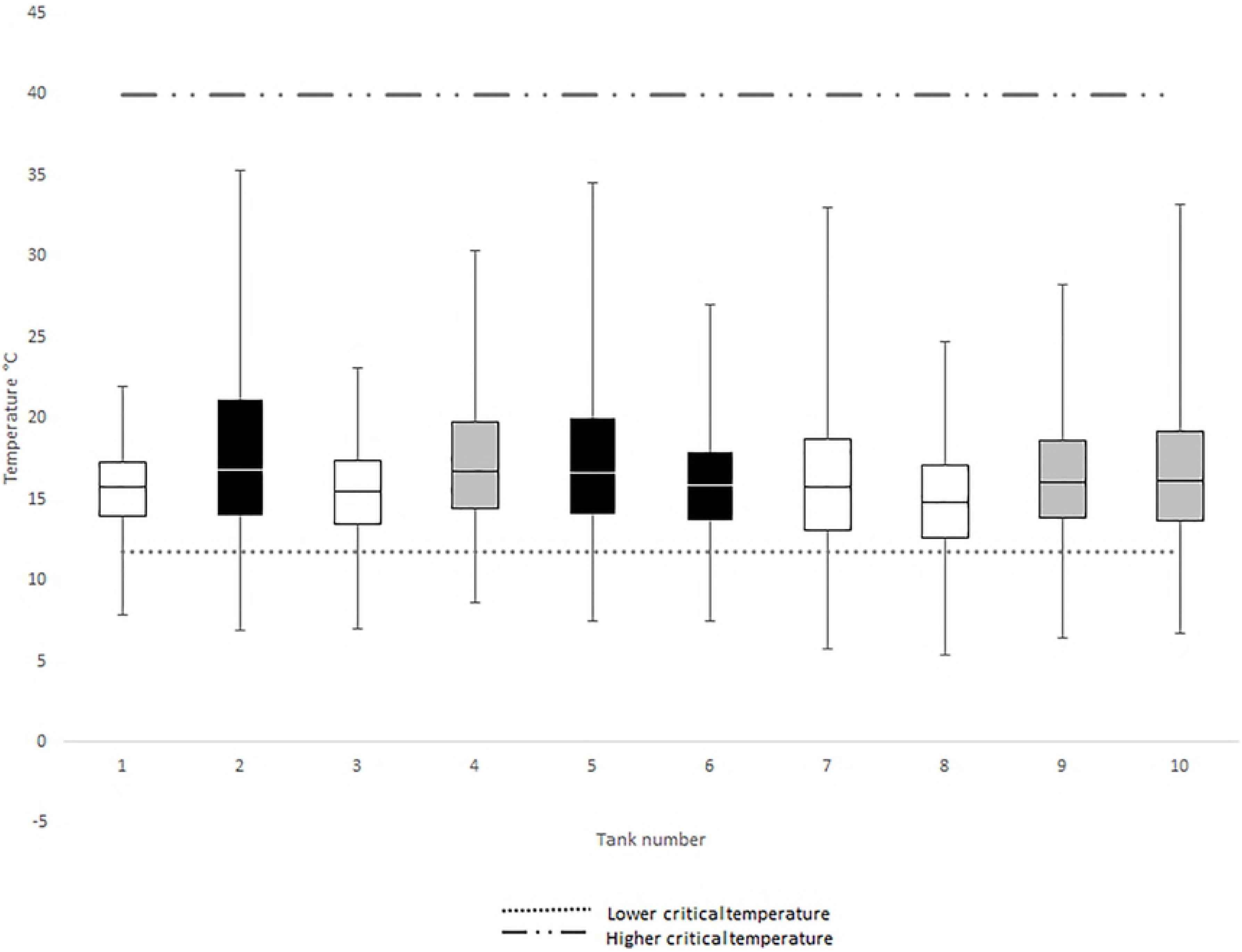
Interquartile range of water temperatures of individual buckets in Brisbane during winter, 2014. Lower critical temperature threshold (11.8°C) for *Aedes aegypti* taken from Eisen et al. (4) and upper threshold (40.0°C) from Richardson et al. (39). Black, grey and white boxes represent low, moderate and high shade conditions, respectively.

Humidity was significantly higher inside rainwater tanks (mean = 78.1, SD = 11.0) than outside (mean = 48.8, SD = 10.7; t(8) = −9.9, *P* < 0.001). Likewise, fortnightly differences between the mean internal (17.2°C, SD = 4.2) and external (9.9°C, SD = 3.7) dew points were significant (t(12) = −4.47, *P* < 0.001). Applying evaporation rates observed in the low shade treatment (assuming a linear relationship over time), we estimate the water in a 9L bucket would take approximately 105 days to evaporate during winter. All abiotic conditions including humidity, dewpoint, salinity, total dissolved solids, changes in volume and evaporation for tanks and buckets are recorded in the Supporting Information (S2 Appendix).

The presence of mosquitoes was observed in tanks fortnightly (Fig 6). *Aedes notoscriptus* was the primary species observed, with a total of 1,820 (mean = 26/ container, SD = 71.64) immature stages counted. Larvae were present in rainwater tanks in 12.5% to 100% of fortnightly surveys (Fig 6). The two tanks that were sealed against any further ingress of rainwater had *Ae. notoscriptus* larvae present only during the first 14 days. The total abundance of immature mosquitoes in first flush devices was 200 (mean = 4.8, SD = 18.7) and larval presence ranged from 12.5% to 62.5% of all surveys (S4 Appendix).

**Fig 6.**
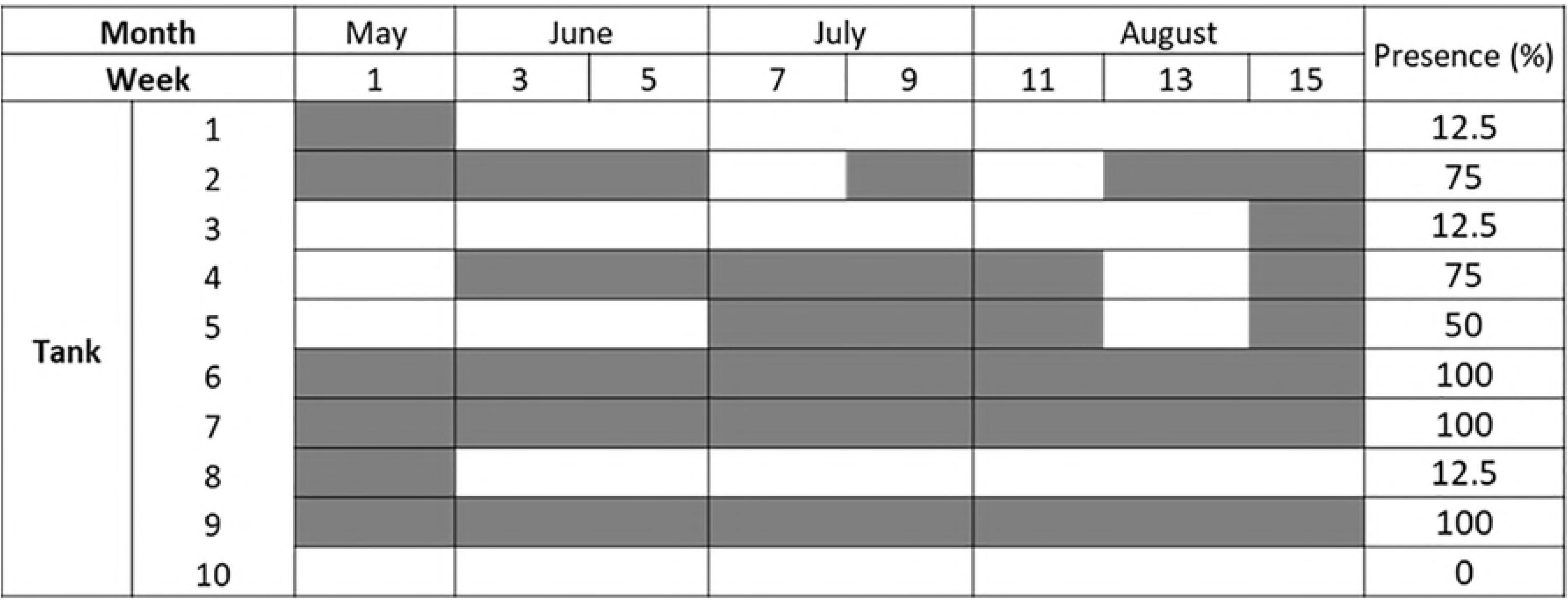
Presence/absence of *Aedes notoscriptus* immatures in sealed rainwater tanks during winter in Brisbane, 2014. Shading represents presence during larval surveys conducted fortnightly. All tanks were sealed, and tanks 1 and 8 had inflows of water removed after the first survey.

### *Aedes aegypti* development and survival under fluctuating temperatures in simulated containers

In environmental chambers, temperatures within containers differed from programmed air temperatures by 1°C (SD = 0.98 tanks, SD = 0.2 buckets). This was due to the thermal capacity of the water stored within the chambers. Rainwater tank temperature simulations increased *Ae. aegypti* survival when compared to bucket simulations (χ^2^= 59.7, df = 1, *P* < 0.001). This was true for tropical strains (Fig 7; S5 Appendix; χ^2^= 18.3, df = 1, *P* < 0.001) and subtropical strains (Fig 8; S5 Appendix; χ^2^= 47.8, df = 1, *P* < 0.001). *Aedes aegypti* from the tropical strain had higher survival in both rainwater tanks (χ^2^=5.2, df = 1, *P* = 0.022) and buckets (χ^2^= 24.7, df = 1, *P* < 0.001) when compared to the subtropical strain.

**Fig 7.**
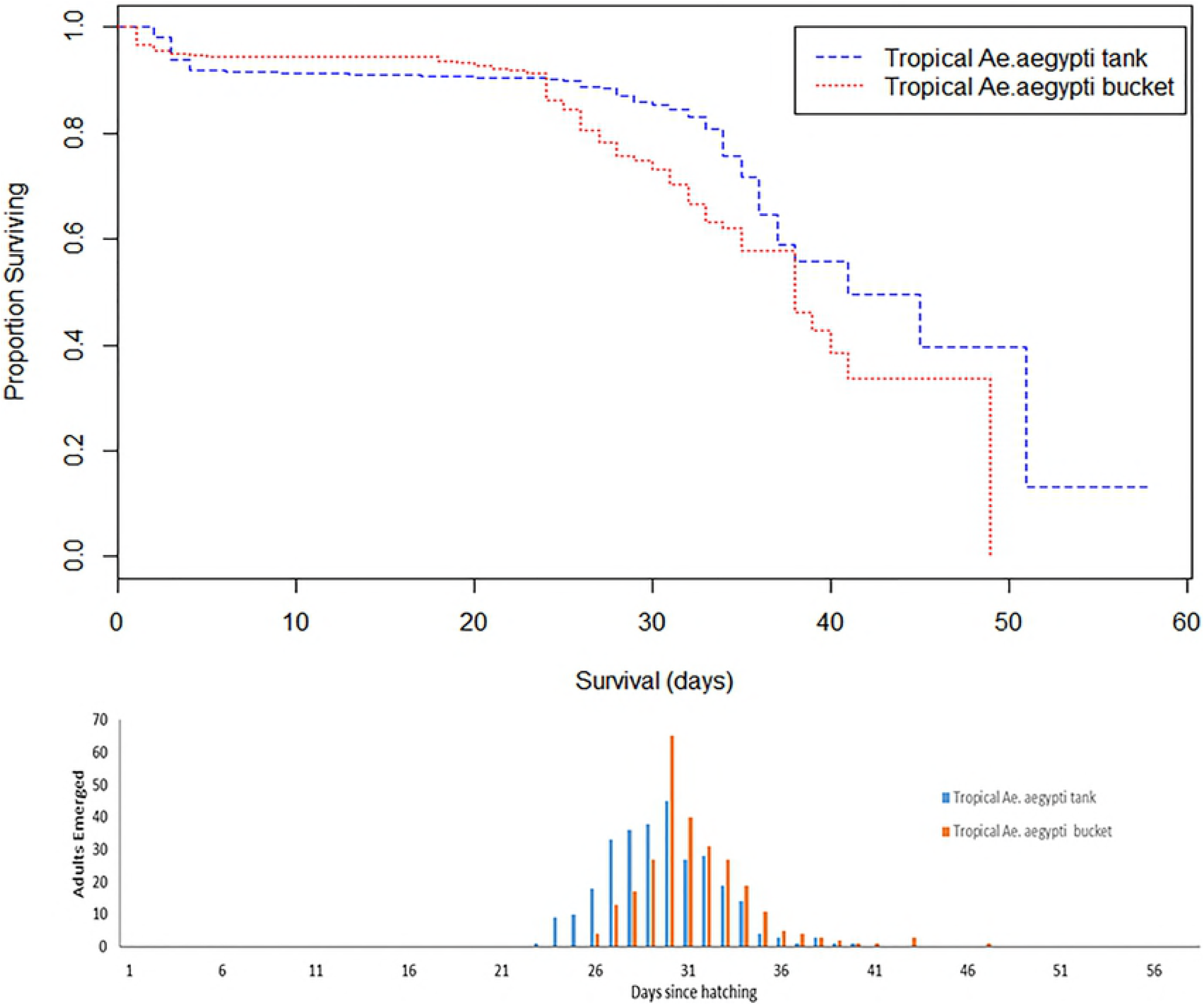
Survival curves and time to emergence of surviving adults comparing tropical *Aedes aegypti* strains in tank and bucket temperature treatments.

**Fig 8.**
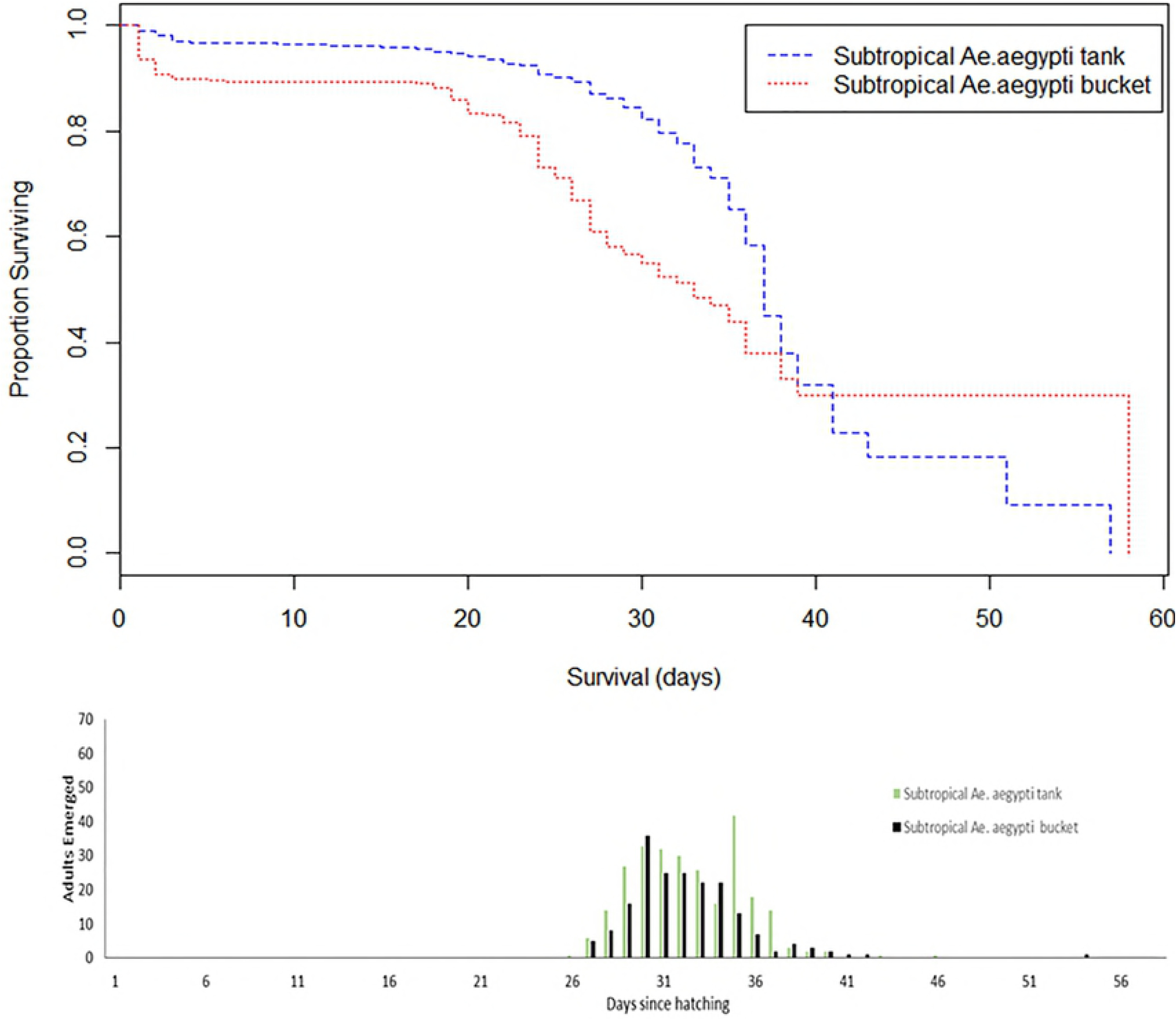
Survival curves and time to emergence of surviving adults comparing subtropical strain of *Aedes aegypti* in tank and bucket temperature treatments.

A comparison of tropical and sub-tropical temperature regimes from tanks showed no differences in mean time to adult emergence (32.5, SE = 0.19; 32.7, SE = 0.20). The same was true for comparisons of subtropical (32.22, SE = 0.23) and tropical (31.37, SE = 0.18) bucket temperatures (F(1,29) = 1.48, *P* = 0.234; S6 Appendix). Analysis indicated that strain and container type had no effect on mean development time (F(1,29) = 0.646, *P* = 0.428), while the interaction effect was not significant (F(1,28) = 0.242, *P* = 0.627).

### Non-linear estimate of *Aedes aegypti* lower critical temperature

We fitted a number of non-linear curves to *Ae. aegypti* development rates and temperatures derived from the published literature (Table 1). Correlations between observed and fitted values were similar across most scenarios. The model with the best correlation that allowed for an estimate of a zero development threshold was the Logan et al., (25) model, which had a correlation of 0.899 (Table 1, Fig 9). This model does not have a parameter for the lower critical threshold, so the equation was solved for the zero development point on the X axis (9.21°C, Table 1, Fig 9). Others estimate this value between 6.55°C and 12.38°C and confidence intervals around these estimates vary considerably (Table 1). Parameter estimates were included for the Sharpe and DeMichele non-linear model (26) traditionally used in simulating *Ae. aegypti* development (27), however, it is impossible to estimate zero development threshold with this model as it never crosses the zero development point on the X axis.

**Table 1.** Fit of non-linear models to *Aedes aegypti* literature. Parameter estimates, estimations of upper and lower critical thresholds, confidence intervals and observed versus expected correlations (Cor) for non-linear models of *Aedes aegypti* development rates under constant and fluctuating temperatures sourced from scientific literature.

| Model (Ref) | devRate Model | Cor | Lower Threshold (95% CI) | Upper Threshold (95% CI) | Parameter 1 | Par 2 | Par 3 | Par 4 | Par 5 | Par 6 |
| --- | --- | --- | --- | --- | --- | --- | --- | --- | --- | --- |
| (28) | kontodimas_04 | 0.894 | 10.84 (1.90) | 41.34 (0.95) | 0.00005 |  |  |  |  |  |
| (29) | perf2_11 | 0.891 | 12.38 (1.95) | 40.67 (0.81) | 0.01349 | 0.193 |  |  |  |  |
| (30) | briere1_99 | 0.883 | 10.00 (3.08) | 40.13 (0.26) | 0.00010 |  |  |  |  |  |
| (30) | briere2_99 | 0.893 | 12.15 (2.55) | 40.44 (0.87) | 0.00007 | 1.439 |  |  |  |  |
| (31) | hilbertLogan_83 | 0.898 | 6.55 (31.3) | 45.38 (197.8) | 3.154 | 62.28 | 7.62 |  |  |  |
| (25) | logan10_76 | 0.899 | 9.21 (NA) | 43.16 (11.2) | 1.128 | 0.135 | 47.22 | 12.644 |  |  |
| (26) | sharpeDeMichel e_77 | 0.900 | N/A | N/A | 32.2 | 14068.5 | -264.96 | -76397 | 282.52 | 87234 |

**Fig 9.**
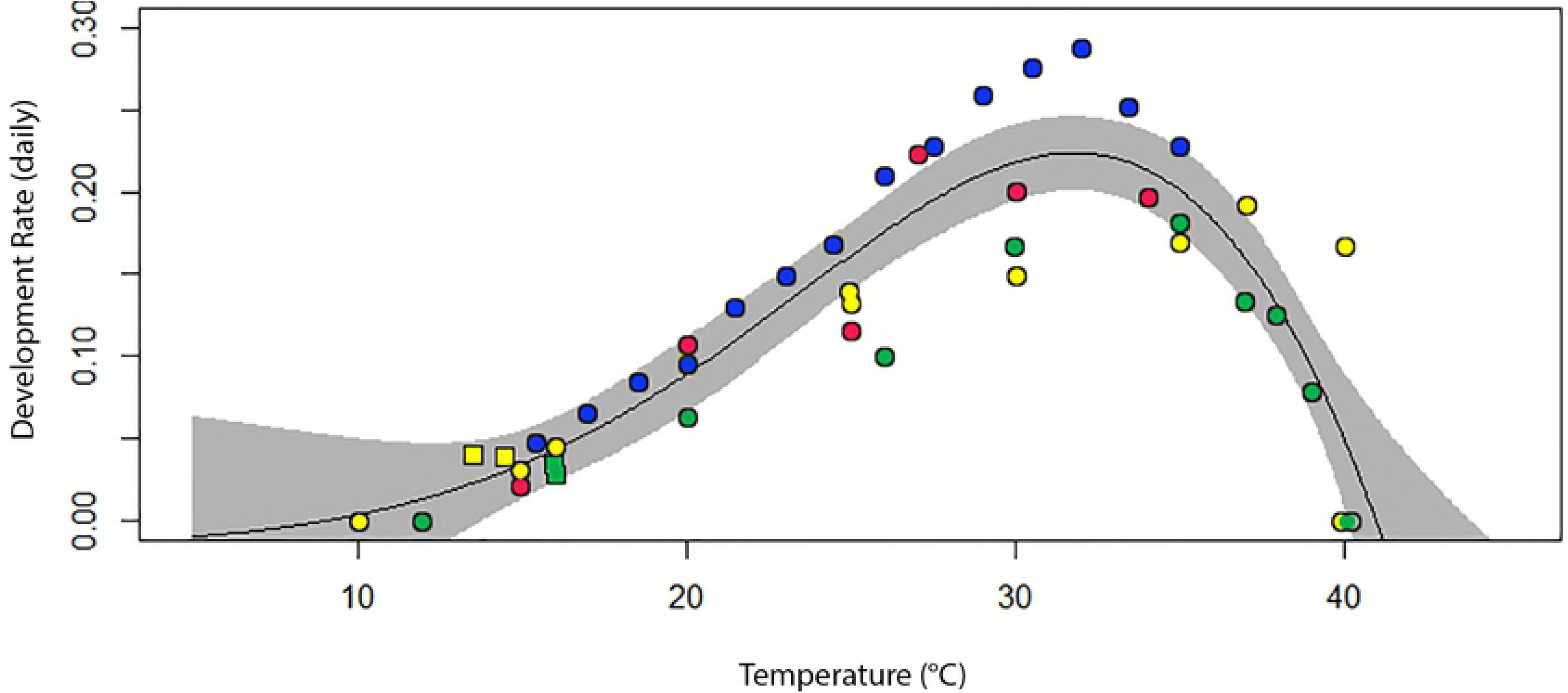
Non-linear development model for *Aedes aegypti.* Comparison of the mean and median time to pupation from the literature, including the current study (13, 38, 39, 42, 43). To these estimates we fit the temperature dependent model developed by Logan et al. (25). Colours represent different continents; Africa (blue), Australia (yellow), South East Asia (green) and North America (red); spots represent constant and squares fluctuating temperatures.

### Degree day estimates of *Aedes aegypti* development

Based on the lower critical temperature of 11.8°C (4, 13), the tropical and subtropical strains of *Ae. aegypti* required 125.7 (SE ± 0.57) and 123.4 (SE ± 0.45) HDDs to develop into adults at a constant 26°C temperature (t(14) = 2.28, *P* = 0.046). There was no significant difference in HDDs required at tank temperatures between the tropical (mean = 80.6, SE ± 2.04) and subtropical strains (mean = 81.1, SE ± 2.16) (t(14) = −0.71, *P* = 0.401). Nor was there any difference in HDDs required for development at bucket temperatures: subtropical (mean = 107.2, SE ± 1.72), tropical (mean = 104.1, SE ± 1.64; t(14) = −1.26, *P* = 0.23). In the bucket treatment, the tropical and subtropical *Ae. aegypti* strains were exposed to 47.2 (SE ± 0.27) and 48.5 (SE ± 0.35) CDD, respectively.

## Discussion

Historically, *Ae. aegypti* was present in Brisbane during the early twentieth century (Cooling when unsealed rainwater tanks and other forms of water storage were common (32-34). Modelling has suggested that conditions in Brisbane are currently inhospitable for the species during winter (2, 35) but the presence of permanent water storage containers, such as rainwater tanks, may change those survival prospects in a subtropical climate.

During winter, tank and bucket water temperatures were comparable, with differences in relative weekly mean temperature differing by less than 2.2°C throughout winter. The largest difference in relative temperature occurred between internal and external air temperatures, with tanks consistently retaining a higher mean air temperature than external air temperatures. Humidity levels of ∼70% and high dewpoints over the surface of the water in rainwater tanks suggests that the air cavity may protect mosquito lifecycle stages when conditions outside are unfavourable. It is likely that these conditions may protect eggs and adults from desiccation stress during periods of low humidity that occur during Australian winters (36, 37).

Our results suggest *Ae. aegypti* can develop and survive throughout winter in Brisbane, in both rainwater tanks and buckets. When *Ae. aegypti* larvae were reared under fluctuating temperature regimes derived from direct observations during the coldest winter month, approximately 50% and 70% survived until adults in buckets and rainwater tanks, respectively. The low thermal inertia exhibited in rainwater tanks resulted in mean weekly minimum and maximum water temperature rarely fluctuating more than ±5°C and seldom breaching the lower critical threshold for development. Rainwater tanks provided a buffered environment, with lower thermal stress, and we suggest that tanks provide a more stable habitat for *Ae. aegypti* larval development when compared to smaller volume containers such as buckets.

No evidence was found to support the hypothesis that *Ae. aegypti* could not survive in simulated winter bucket temperatures from Brisbane. Instead, 48-67% of larvae were observed to survive in bucket treatments representing the coldest week observed, where the mean temperature (13.5°C) was close to the lower critical temperature (4). This result contradicts modelling by Kearney et al. (2) and Williams et al. (35). However, conditions in our study had consistent volumes and ideal nutrition and water quality, so results should be interpreted as optimal conditions for survival within the temperatures tested. The dark colouring of buckets likely lowered solar reflectance, therefore enhancing absorption of solar radiation and increasing the upper fluctuation into the optimal temperature range. However, during the night buckets of all shade regimes stabilized with air temperature around 12am at time resulting in minimum temperatures dropping below the lower critical threshold and we observed larvae capable of surviving temperatures down to 4.5°C for short periods of time.

Fluctuating temperatures can have varying effects on life span, particularly due to the length of time and amplitude of exposure outside of optimal conditions (14). This suggests that the higher survival observed in the tank treatments (67% - 76%) was the result of lower cold stress (CDD) when compared with bucket treatments. Low temperature stresses may account for differences between our survival results and those contrasting result of Carrington et al. (13), who observed a higher survival in their large fluctuation treatment. Temperature fluctuations in the current study had lower maximum and minimum mean temperatures (22.8°C and 4.5°C respectively for the large fluctuation) which may have resulted in larvae experiencing longer periods below the lower critical threshold when compared with Carrington et al. (13; range 25.3°C to 6.7°C). Furthermore, our study did not compare treatments with similar means due to differences in volumes observed in field containers and on which we based our temperature models. Our differences in survival support the hypothesis that rainwater tanks (small fluctuation) can provide a thermally buffered habitat during periods of stress, and are likely to increase survival of immatures to adult emergence when compared to containers of smaller volume.

Although there were differences in survival between strains, results for temperature regimes in our study were above those observed in previous studies where constant temperatures were applied (13, 38, 39). Previous studies sourced their strain from a similar location in North Queensland (38, 39) so it is unlikely that variations observed in the current study were attributable to population differences. Therefore, we suggest that fluctuating temperatures, whether high or low, can increase the survival of *Ae. aegypti* around lower thresholds. Higher survival in the tropical strain suggests that results do not support the hypothesis that southern populations are adapted to colder temperatures. As far as we are aware there are no records of temperature adaptation in *Ae. aegypti* so this is perhaps unsurprising. As adaptation is facilitated by genetic diversity, driven by the accumulation of beneficial mutations, it is also possible that southerly Queensland populations may have very limited genetic diversity (40) and therefore limited scope to adapt.

The lower critical temperature for *Ae. aegypti* is clearly less than what is currently accepted as the value for this limit (11.78 ∼11.8°C) which is typically calculated using linear regression to estimate the point where the function crosses the temperature-axis (4, 13, 38). Estimating the lower critical temperature via constant temperatures is highly artificial (15). It may be that the fluctuating temperature studies we designed also give more accurate empirical thresholds. In reality, development rates at the lower threshold tend to decay in an exponential fashion and thermal minimums estimated using linear functions will have a high margin of error compared to estimations using non-linear methods (15). Thus our findings, which indicated a threshold around 10°C, suggest that the lower critical threshold for *Ae. aegypti* is likely lower than previous estimates and studies (particularly those using degree days) may have underestimated the ability of the species to endure colder temperatures.

Our estimates of development time in colder temperatures were consistent with other studies of Australian *Ae. aegypti* populations (38, 39). These studies estimated mean emergence times of 30-39 days for constant temperatures regimes of 15°C and 16°C (38, 39). Interestingly, no significant differences in mean development time were observed between the fluctuating temperature regimes in the current study. However, we did observe differences in the number of HDDs required for development when applying the traditional lower critical temperature, with mosquitoes under the tank regime requiring fewer degree days than the bucket regime. It is likely that HDD estimates will be inaccurate when assuming the 11.8°C threshold. For example, the number of HDDs calculated for a small fluctuation around an inaccurate lower critical temperature, will tend to underestimate the total HDDs required for development compared to rates entering the linear part of the developmental curve (15). Thus, interpreting development rates that apply linear relationships around thermal minimums must be done with caution.

*Aedes notoscriptus* larvae, the native container inhabiting species, were consistently present in sealed tanks throughout the winter. This suggests that roof guttering or piping may play an important role in ‘seeding’ tanks with larval mosquitoes during rainfall events. The role that gutters play as a source of container inhabiting mosquitoes has been identified previously in north Queensland (41). In our study, gutters with high levels of organic matter or sitting water were likely responsible for eggs or larvae being washed into tanks during frequent rainfall events (S7 Appendix). When tanks are sealed, there is little chance of adult mosquitoes escaping but any small gap in the multiple seals, mesh openings and plastic covers typical of rainwater tanks, would convert them into highly productive habitats. Furthermore, the presence of *Ae. notoscriptus* larvae was observed in first flush devices throughout winter, suggesting that attendant infrastructure such as these containers, could seed larvae into tanks and contribute to local mosquito populations.

Our findings have important implications for estimating the potential distribution of *Ae. aegypti* and demonstrating the risk of re-establishment in southern Queensland where larval habitat is readily available in the form of rainwater tanks (8). Considering the historical presence of *Ae. aegypti* as far south as the Victorian boarder, it is no surprise that larvae are capable of surviving winter temperatures in Brisbane. Kearney et al. (2) and Richardson et al. (39) postulated that rainfall was not sufficient in Brisbane for small containers, such as buckets, to act as larval habitat for *Ae. aegypti* throughout the year. However, field observations indicated that, in winter any container holding >3L or that retains water for >32 days is potentially productive during even the coldest months. Buckets represent one of the most common and productive containers for urban mosquitoes in Brisbane (8).

We conclude that rainwater tanks and buckets provide ideal larval habitat for the (re)establishment and persistence of invasive mosquito species in areas where low rainfall and temperatures might make establishment difficult. These containers ensure that sub-optimal landscapes can be unwittingly manipulated by human behaviours to support the establishment of invasive disease vectors across new urban areas. If rainwater tanks and other key containers are not managed appropriately, large areas of southern Australia may see the return of *Ae. aegypti* with tremendous implications for public health and the management of imported cases of dengue, chikungunya, Zika and yellow fever.

## Acknowledgements

We would like to thank the tank owners, Miriam and Rob Gibson, Saxon and Rowena Gamble, Camilla and Rob Burford, Anne Bourne, Jason and Sonia Morgan, Iain Crawford and Charl and Karin Taylor for allowing us to access their premises. Vehicular assistance from the Mosquito and Arbovirus Research Committee. The Cairns Public Health Unit and Greg Crisp and Jason Gilmore from the Wide Bay Public Health Unit for assistance in collecting eggs for each mosquito colony. Anna Marcora for assisting in field temperature collections. Funding was supplied through an Integrated National Resources and Management grant from the University of Queensland and the Commonwealth Science and Industrial Research Organisation. The authors have no conflict of interests to declare.

## Author Contributions

BT, JD, GD and MZ conceived the ideas and designed methodology;

BT and JD collected the data;

BT analysed the data;

BT, JD, MZ, NS, CJ and GD contributed to the writing of the manuscript.

## Appendix

**S1. Temperature Regimes.** Environmental chamber temperatures used to determine survival of *Aedes aegypti* in different container categories from Brisbane, Australia.

**S2. Rainwater Tank Conditions.** Abiotic conditions within rainwater tanks and buckets in Brisbane during winter (July) 2014. Levels represent 0-32% shade cover (1), 33-65% (2) and 66-100% (3).

**S3. Weather Records.** Temperatures recorded in air at Archerfield Airport, (−27.57o S, 153.01o E), tanks and buckets from Brisbane during winter (1st June until 31st August), 2014.

**S4. Mosquito Presence in Rainwater Tanks.** Presence of *Aedes notoscriptus* immatures (grey shading) in first flush devices during winter in Brisbane, 2014. Numbers correspond to the tank which contained devices. For example, tank 10 had two separate first flush devices (10a,10b) on downpipes entering tank. Volume measures the mean volume found in each device throughout the field survey. Presence represents the percentage of surveys where at least one *Ae. notoscriptus* immature was sampled from the device.

**S5. *Aedes aegypti* Survival.** Survival of tropical and subtropical *Aedes aegypti* strains in rainwater tank (small fluctuation), buckets (large fluctuation) and 26°C control (constant) treatments.

**S6. *Aedes aegypti* Survival.** Mean, standard error, minimum and maximum development time for tropical and subtropical *Aedes aegypti* strains in rainwater tank (small fluctuation), buckets (large fluctuation) and 26°C control (constant) treatments.

**S7. Productive Infrastructure.** Roof gutters observed that likely increased productivity of rainwater tanks during winter in Brisbane, 2014.

## References

1. Kearney M, Porter W. Mechanistic niche modelling: combining physiological and spatial data to predict species’ ranges. Ecol Lett. 2009;12(4):334–50. doi:10.1111/j.1461-0248.2008.01277.x

2. Kearney M, Porter WP, Williams C, Ritchie S, Hoffmann A. Integrating biophysical models and evolutionary theory to predict climatic impacts on species’ ranges: the dengue mosquito *Aedes aegypti* in Australia. Func Ecol. 2009;23(3):528–38. doi:10.1111/j.1365-2435.2008.01538.x

3. Russell RC, Currie BJ, Lindsay MD, Mackenzie JS, Ritchie SA, Whelan PI. Dengue and climate change in Australia: predictions for the future should incorporate knowledge from the past. Med J Aust. 2009;190(5):265–8.

4. Eisen L, Monaghan AJ, Lozano-Fuentes S, Steinhoff DF, Hayden MH, Bieringer PE. The impact of temperature on the bionomics of *Aedes (Stegomyia) aegypti*, with special reference to the cool geographic range margins. J Med Entomol. 2014;51(3):496–516. doi:10.1603/ME13214

5. Jetten TH, Focks DA. Potential changes in the distribution of dengue transmission under climate warming. Am J Trop Med Hyg. 1997;57(3):285–97. doi:10.4269/ajtmh.1997.57.285

6. Bader CA, Williams CR. Mating, ovariole number and sperm production of the dengue vector mosquito *Aedes aegypti* (L.) in Australia: broad thermal optima provide the capacity for survival in a changing climate. Physiol Entomol. 2012;37(2):136–44. doi:10.1111/j.1365-3032.2011.00818.x

7. Beebe NW, Cooper RD, Mottram P, Sweeney AW. Australia’s dengue risk driven by human adaptation to climate change. PLoS NTD. 2009;3(5):e429. doi:10.1371/journal.pntd.0000429

8. Trewin BJ, Kay BH, Darbro JM, Hurst TP. Increased container-breeding mosquito risk owing to drought-induced changes in water harvesting and storage in Brisbane, Australia. Int Health. 2013;5(4):251–8. doi:10.1093/inthealth/iht023

9. Jansen CC, Beebe NW. The dengue vector *Aedes aegypti*: what comes next. Microbes Infect. 2010;12(4):272–9. doi:10.1016/j.micinf.2009.12.011

10. Lumley GF, Taylor FH. Dengue: School of Public Health and Tropical Medicine (University of Sydney); 1943.

11. Russell RC. Mosquito-borne arboviruses in Australia: the current scene and implications of climate change for human health. Int J Parasitol. 1998;28(6):955–69. doi:10.1016/S0020-7519(98)00053-8

12. Maréchal Y. The hydrogen bond and the water molecule: The physics and chemistry of water, aqueous and bio-media: Elsevier; 2006.

13. Carrington LB, Armijos MV, Lambrechts L, Barker CM, Scott TW. Effects of fluctuating daily temperatures at critical thermal extremes on *Aedes aegypti* life-history traits. PLoS One. 2013;8(3):e58824. doi:10.1371/journal.pone.0058824

14. Colinet H, Sinclair BJ, Vernon P, Renault D. Insects in fluctuating thermal environments. Ann Rev Entomol. 2015;60:123–40. doi:10.1146/annurev-ento-010814-021017

15. Allsopp PG, Daglish GJ, Taylor MFJ, Gregg PC, Zalucki MP. Measuring development of *Heliothis* species. Heliothis: research methods and prospects. 1991:90-101. doi:10.1007/978-1-4612-3016-8_8

16. Ruel JJ, Ayres MP. Jensen’s inequality predicts effects of environmental variation. Trends Ecol Evol. 1999;14(9):361–6. doi:10.1016/S0169-5347(99)01664-X

17. Bureau of Meteorology. Climate Data Online: Bureau of Meteorology; 2015 [Available from: http://www.bom.gov.au/climate/data/.

18. ESRI World Imagery, cartographer World Topographic Map2018. “Topographic” [basemap]. “World Topographic Map”. November 7, 2018. http://www.arcgis.com/home/item.html?id=30e5fe3149c34df1ba922e6f5bbf808f. Accessed November 19, 2018.

19. Knox TB, Yen NT, Nam VS, Gatton ML, Kay BH, Ryan PA. Critical Evaluation of Quantitative Sampling Methods for *Aedes aegypti* (Diptera: *Culicidae*) Immatures in Water Storage Containers in Vietnam. J Med Entomol. 2007;44(2):192–204. doi:10.1093/jmedent/44.2.192

20. Rasic G, Endersby NM, Williams C, Hoffmann AA. Using *Wolbachia*-based release for suppression of *Aedes* mosquitoes: insights from genetic data and population simulations. Ecol Appl. 2014;24(5):1226–34. doi:10.1890/13-1305.1

21. Ulrich JN, Beier JC, Devine GJ, Hugo LE. Heat Sensitivity of wMel *Wolbachia* during Aedes aegypti Development. PLoS NTD. 2016;10(7):e0004873. doi:10.1371/journal.pntd.0004873

22. Hugo LE, Kay BH, O’neill SL, Ryan PA. Investigation of environmental influences on a transcriptional assay for the prediction of age of *Aedes aegypti* (Diptera: Culicidae) mosquitoes. J Med Entomol. 2010;47(6):1044–52. doi:10.1603/ME10030

23. Pepe MS, Fleming TR. Weighted Kaplan-Meier statistics: a class of distance tests for censored survival data. ISO4. 1989:497–507.

24. R Core Team. R: A language and environment for statistical computing.2018; (3.5.1). Available from: http://www.R-project.org/.

25. Logan JA, Wollkind DJ, Hoyt SC, Tanigoshi LK. An Analytic Model for Description of Temperature Dependent Rate Phenomena in Arthropods 1. Enviro Entomol. 1976;5(6):1133–40. doi:10.1093/ee/5.6.1133

26. Sharpe PJ, DeMichele DW. Reaction kinetics of poikilotherm development. J Theor Biol. 1977;64(4):649–70. doi:10.1016/0022-5193(77)90265-X

27. Focks DA, Daniels E, Haile DG, Keesling JE. A simulation model of the epidemiology of urban dengue fever: literature analysis, model development, preliminary validation, and samples of simulation results. Am J Trop Med Hyg. 1995;53(5):489–506. doi:10.4269/ajtmh.1995.53.489

28. Kontodimas DC, Eliopoulos PA, Stathas GJ, Economou LP. Comparative Temperature-Dependent Development of *Nephus includens* (Kirsch) and *Nephus bisignatus* (Boheman) (Coleoptera: *Coccinellidae*) Preying on *Planococcus citri* (Risso) (Homoptera: *Pseudococcidae*): Evaluation of a Linear and Various Nonlinear Models Using Specific Criteria. Enviro Entomol. 2004;33(1):1–11. doi:10.1603/0046-225X-33.1.1

29. Shi P, Ge F, Sun Y, Chen C. A simple model for describing the effect of temperature on insect developmental rate. J Asia Pac Entomol. 2011;14(1):15–20. doi:10.1016/j.aspen.2010.11.008

30. Briere J-F, Pracros P, Le Roux A-Y, Pierre J-S. A Novel Rate Model of Temperature-Dependent Development for Arthropods. Enviro Entomol. 1999;28(1):22–9. doi:10.1093/ee/28.1.22

31. Hilbert DW, Logan JA. Empirical Model of Nymphal Development for the Migratory Grasshopper, *Melanoplus sanguinipes* (Orthoptera: *Acrididae*) 1. Enviro Entomol. 1983;12(1):1–5. doi:10.1093/ee/12.1.1

32. Cooling LE. Report on Mosquito Survey of the Brisbane Metropolitan Area, 1923. Australia Dept Health. 1923.

33. Hamlyn-Harris R. The elimination of *Aedes argenteus* Poiret as a factor in dengue control in Queensland: Liverpool School of Tropical Medicine; 1931.

34. Trewin BJ, Darbro JM, Jansen CC, Schellhorn NA, Zalucki MP, Hurst TP, et al. The elimination of the dengue vector, *Aedes aegypti*, from Brisbane, Australia: The role of surveillance, larval habitat removal and policy. PLoS NTD. 2017;11(8):e0005848. doi:10.1371/journal.pntd.0005848

35. Williams CR, Bader CA, Kearney MR, Ritchie SA, Russell RC. The extinction of dengue through natural vulnerability of its vectors. PLoS NTD. 2010;4(12):e922. doi:10.1371/journal.pntd.0000922

36. Russell B, Kay B, Shipton W. Survival of *Aedes aegypti* (Diptera: *Culicidae*) eggs in surface and subterranean breeding sites during the northern Queensland dry season. J Med Ent. 2001;38(3):441–5. doi:10.1603/0022-2585-38.3.441

37. Faull KJ, Williams CR. Intraspecific variation in desiccation survival time of *Aedes aegypti* (L.) mosquito eggs of Australian origin. J Vect Ecol. 2015;40(2):292–300. doi:10.1111/jvec.12167

38. Tun-Lin W, Burkot TR, Kay BH. Effects of temperature and larval diet on development rates and survival of the dengue vector *Aedes aegypti* in north Queensland, Australia. Med Vet Ent. 2000;14(1):31–7. doi:10.1046/j.1365-2915.2000.00207.x

39. Richardson K, Hoffmann AA, Johnson P, Ritchie S, Kearney MR. Thermal Sensitivity of *Aedes aegypti* From Australia: Empirical Data and Prediction of Effects on Distribution. J Med Ent. 2011;48(4):914–23. doi:10.1603/me10204

40. Endersby-Harshman NM. *Aedes aegypti* from Gin Gin, Queensland – potential for a successful suppression/extinction Wolbachia program based on population genetic data. 2014.

41. Montgomery BL, Ritchie SA. Roof gutters: a key container for *Aedes aegypti* and *Ochlerotatus notoscriptus* (Diptera: *Culicidae*) in Australia. Am J Trop Med Hyg. 2002;67(3):244–6. doi:10.4269/ajtmh.2002.67.244

42. Gilpin ME, McClelland GA. Systems analysis of the yellow fever mosquito *Aedes aegypti*. Fortschr Zool. 1979;25(2-3):355.

43. Rueda LM, Patel KJ, Axtell RC, Stinner RE. Temperature-Dependent Development and Survival Rates of *Culex quinquefasciatus* and *Aedes aegypti* (Diptera: *Culicidae*). J Med Ent. 1990;27(5):892–8. doi:10.1093/jmedent/27.5.892

